# Expression of *dlx* genes in the normal and regenerating brain of adult zebrafish

**DOI:** 10.1101/2020.02.11.943605

**Authors:** Hellen Weinschutz Mendes, Mariam Taktek, Thomas Duret, Marc Ekker

## Abstract

Dysfunctions in the GABAergic system lead to various pathological conditions and impaired inhibitory function is one of the causes behind neuropathies characterized by neuronal hyper excitability. The *Dlx* homeobox genes are involved in the development of nervous system, neural crest, brachial arches and developing appendages. *Dlx* genes also take part in neuronal migration and differentiation during development, more precisely, in the migration and differentiation of GABAergic neurons. Functional analysis of *dlx* genes has mainly been carried out in developing zebrafish embryos and larvae; however information regarding the expression and roles of these genes in the adult zebrafish brain is still lacking. The extensive neurogenesis that takes place in the brain of adult zebrafish makes them a good model for the visualization of mechanisms involving *dlx* genes during adulthood in physiological conditions and during regeneration of the nervous system. We have identified the adult brain regions where transcripts of *dlx1a, dlx2a, dlx5a* and *dlx6a* genes are normally found and have confirmed that within telencephalic domains, there is high overlapping expression of the four *dlx* paralogs with a marker for GABAergic neurons. Co-localization analyses carried with the Tg(*dlx6a*-1.4kb*dlx5a/dlx6a*:GFP) reporter line have also shown that in some areas of the diencephalon, cells expressing the *dlx5a/6a* bigene may have a neural stem cell identity by co-localizing with a Sox2 antibody. Furthermore, investigations in a response to stab wound lesions, have demonstrated a possible participation of the *dlx5a/6a* bigene, most likely, of *dlx5a* during the regeneration of the adult zebrafish brain. These data suggest a possible participation of *dlx*-expressing cells during brain regeneration in adult zebrafish and also provide information on the role of *dlx* genes under normal physiological conditions in adults.

## Introduction

The transcription factors encoded by *Dlx* genes play key roles in the patterning of the vertebrate limb and the central nervous system (CNS) (1), more specifically, *Dlx* genes are required in the development of the mammalian brain (2,3). These genes are required for correct migration and differentiation of progenitors that will later give rise to GABAergic interneurons (4). *Dlx1*^-/-^*/Dlx2*^*-/-*^ mutant mice show a loss of GABAergic interneuron differentiation in the ventral telencephalon, supporting the notion of this requirement for *Dlx* genes in the differentiation to GABAergic interneurons (5). In the case of zebrafish, *dlx* genes are also expressed in the developing brain (6) (7). At 24-48 hours-post-fertilization (hpf), there is partial overlapping expression of *dlx* and *gad1* genes in the zebrafish forebrain (8). Nevertheless, information regarding the activity and functions of *Dlx* genes in the adult brain is still scarce and non-existent in the zebrafish.

The majority of vertebrates have six *Dlx* genes which are organized in convergently transcribed bigene pairs, namely *Dlx1/Dlx2, Dlx3/Dlx4* and *Dlx5/Dlx6* (9) (10) (11). In zebrafish, the *dlx1a/dlx2a* and *dlx5a/dlx6a* (orthologs of mouse *Dlx1/Dlx2* and *Dlx5/Dlx6*) are expressed in the developing brain. Previously described cis-regulatory regions are essential for driving the expression of these bigenes in the brain. The deletion of one of these regions in mice, I56ii, has been shown to impair the expression of *Dlx* genes and of potential targets including *Gad2* and other striatal markers (12). The identification of such regulatory elements was a starting point for the generation of the Tg(*dlx6a*-1.4kb*dlx5a/dlx6a*:GFP) reporter line that mimics the endogenous expression patterns of *dlx5a/dlx6a* genes in the forebrain (11,13).

Using the Cre/LoxP system for lineage tracing, Solek and collaborators have reported a detailed analysis of fate decisions for *dlx1a/2a*- and *dlx5a/6a-*expressing cells. In some instances, labeling larval *dlx5a/6a*-expressing cells, but not *dlx1a/2a*-expressing cells, have resulted in massively expanding, widespread clonal expansion throughout the adult brain (14). These analyses provided some indications concerning the role of the progeny of *dlx*-expressing cells in the adult zebrafish brain, but further analyses are necessary for investigating these *dlx*-expressing cells in the adult brain.

Different investigations have implicated the *dlx* genes in a group of complex genetic regulatory networks responsible for proper establishment of neuronal diversity in the CNS during development. Interestingly, the establishment of new neurons also takes place in the adult zebrafish brain where multiple areas present constitutive proliferation (15,16). In several mammals and bird species, constitutive active neurogenic domains are restricted to the forebrain, whereas in the zebrafish, new neurons are generated in most brain regions throughout adult life (reviewed (17,18)). High rates of adult neurogenesis are present in thirteen to sixteen distinct neural stem cell niches along the adult zebrafish brain. The adult zebrafish brain possesses regeneration capability, which makes this animal an ideal model to study the mechanisms involved in brain regeneration and the different genes participating in regeneration responses within the CNS (19) (reviewed in (20,21)). Therefore, a remarkable capacity to regenerate the CNS following mechanical or chemical insult is present in the zebrafish (17).

The important roles of *dlx* genes during the development and specification of GABAergic neurons and the potential for regenerative investigations of adult zebrafish led us to carry investigations on *dlx* paralogs in the adult zebrafish brain. In this work, we report expression of all four *dlx* paralogs in the adult zebrafish brain and show that the majority of cells expressing these genes are in fact GABAergic neurons. Using the previously described transgenic line Tg(*dlx6a*-1.4kb*dlx5a/dlx6a*:GFP) we carried co-localization analyses which revealed that *dlx5a/6a*-expressing cells present a neural stem cell identity in specific areas of the forebrain and midbrain of adult zebrafish. Furthermore, during a regeneration response following stab injury, we observed an up-regulation of *dlx5a* and of the *dlx5a/6a* bigene.

## Materials and Methods

### Animal care, husbandry and strains

All experiments were conducted using protocols approved by the University of Ottawa Animal Care Committee. Adult zebrafish were housed in circulating water at 28.5C and 14-h light cycle, following standard procedures previously described (Westerfield, 2000). Zebrafish embryos were obtained from the natural spawning of adult zebrafish. Facility-raised wild type and adult zebrafish of a reporter line, Tg(*dlx6a*-1.4kb*dlx5a/dlx6a*:GFP), were used. In this reporter line, the green fluorescent protein (GFP) is expressed under the control of *cis*-regulatory elements of the *dlx5a/6a* bigene, namely the I56i and I56ii intergenic region and a 3.5kb fragment of the *dlx6a* 5’-flanking region (9) (11). In this line the GFP reporter recapitulates the expression of *dlx5a* and/or *dlx6a* (9,22,23).

### Brain lesions and collection of zebrafish brain tissue

Surgeries were performed on adult zebrafish raging from 3mpf to 1ypf as described by Schmidt (2014) (24). Briefly, after being anesthetized with a 0.4% Tricaine solution, fish are injured by the insertion of sterile 30g needle directly and vertically trough the skull into the medial region of one telencephalic hemisphere under a dissecting microscope (24). Control specimens are anesthetized, but no injury is provoked. After surgery, adult zebrafish were maintained under the same conditions as the rest of the colony. Lesion and control specimens were euthanized at 3 or 7 days post-lesion (dpl) for analyses.

Fish were euthanized by immersion in Tricaine MS-222 solution at 0.8% in system water, the upper skull was removed and the whole head was fixed in 4% PFA/PBS overnight at 4 °C. After whole brain was dissected, they were post fixed for additional 30 min. The tissue was then washed several times with PBS and equilibrated with 30% sucrose/PBS overnight at 4 °C. Whole brains were then incubated in a solution of 1:2 30% Sucrose:OCT Compound (Tissue-Tek, VWR Canada) for 30 min, placed in cryomolds and frozen in liquid nitrogen. Cryosections of 14-16 µm were obtained with a CM3050S cryostat (Leica, Concord, ON) in duplicate, triplicate or quadruplicate slides.

### Immunohistochemistry (IHC) and double IHC

Sections were first rehydrated in PBST (PBS with 0.1% Tween-20), and blocked in 10% fetal bovine serum in PBST for at least 2 hours at room temperature. The primary antibodies were used at different dilutions according to the manufacturer’s instructions and optimization of the protocol (Table 1). The primary antibody incubation was carried out overnight at 4°C in 1% fetal bovine serum in PBST. Sections were then washed 3 times /15 min with PBST and incubated with the secondary antibodies for 2 h at room temperature (Table 1). Sections were again washed with PBST and nuclei visualized with DAPI (Life Technologies, Burlington, ON). The Calbindin, Calretinin, PCNA and TH antibodies required an extra step of antigen retrieval. Sections were treated for 20 min at 85 °C in 0.01 M sodium citrate/0.05% Tween-20 solution and cooled down to RT for 15 minutes prior to blocking. Images were acquired with either a Nikon A1 Confocal microscope or a Zeiss AxioPhot Fluorescence Microscope and treated with NIS-Elements Advanced Research Software or ImageJ.

**Table 1.** Co-localization of *dlx* genes and different markers.

| Brain areas/ Markers | gad65a<sup>+</sup> | Calbindin<sup>*</sup> | Calretinin<sup>*</sup> | TH<sup>*</sup> | Sox2<sup>*</sup> | GFAP<sup>*</sup> | PCNA<sup>*</sup> |
| --- | --- | --- | --- | --- | --- | --- | --- |
| Ventral Telencephalon | +++ | + | - | + | - | + | + |
| Dorsal Telencephalon | +++ | + | - | - | + | - | - |
| Preoptic Area | ++ | ++ | + | - | +++ | - | - |
| Periventricular gray zone | n.c. | + | + | - | + | - | - |
| Caudal zone of Hypothalamus | + | - | - | + | +++ | - | - |
| Dorsal zone of Hypothalamus | + | - | - | - | - | - | - |

n.c.: Not conclusive
- no co-localization observed
+ very scarce co-localization (1 – 15% of cells present co-localization)
++ 15% to 50% of cells present co-localization
+++ Over 50% of cells present co-localization

### Riboprobes and In-situ hybridization (ISH)

Expression of *dlx1a, dlx2a, dlx5a, dlx6a* and *gad65* was determined using In situ hybridization assays with antisense mRNA probes on crysections as described (Dorsky et al., 1995). Antisense mRNA probes were labeled with digoxygenin-dNTP or dinitrophenol DNP-11-UTP and synthesized from cDNA clones, *dlx1a* (Ellies et al., 1997), *dlx2a* and *dlx5a* (Akimenko et al., 1994), *dlx6a* (Ellies et al., 1997) and *gad65* (Martin 1998). Vectors containing the cDNA clones were linearized with *Bam*HI, *Eco*RI or *Xho*l and the antisense riboprobes were synthesized using either the T7 or T3 polymerase as required.

Brain sections, stored at −20°C, were thawed at room temperature for 30 minutes before the experiment. Hybridization was carried out overnight at 70°C in a humidified chamber. Slides were washed twice with Solution A (50% Formamide, 5% 20x SSC in dH_2_0) and twice with TBS. Blocking with 10% FBS TBST was carried for 2 hours in RT. Detection of hybridized probes was performed with anti-DIG antibodies AP fragments (Roche, Basel Switzerland; dilution 1:1000) overnight at 4°C. After four TBST washes, staining was developed with NBT/BCIP for 6–18h (Sigma, St-Louis, MO). Images were acquired with a Zeiss AxioPhot Fluorescence Microscope.

### Double Fluorescent In-situ Hybridization (dblFISH)

Sections were treated with 2% H_2_O_2_ in PBS to inactivate endogenous peroxidase followed by incubation with anti-DIG antibodies POD fragments in combination with anti-DNP POD (Roche, Basel Switzerland; dilution 1:1000). Incubations with these antibodies were done separately at 4°C, overnight, for each of the antibodies. Staining with tyramide Cy3 solution or Fluorescein in PBS/Tween (1:100) was carried for 10 min each (Perkin-Elmer, Woodbridge, Ontario). Images were acquired with a Nikon A1 confocal microscope and/or Zeiss AxioPhot Fluorescence Microscope and treated with NIS-Elements Advanced Research Software or ImageJ.

### RNA extraction, cDNA synthesis and qRT-PCR

Quantification of *dlx1a, dlx2a, dlx5a* and *dlx6a* RNA transcripts within brain tissues, was performed on a BioRad CFX96 quantative Reverse Transcription PCR detection system using SsoFast EvaGreen (BioRad) fluorescent dye supermix and specific primers for each gene (Supp Table 1). Primers were designed in separate exon sequences using NCBI’s Primer-BLAST Program (Primer-Blast, National Center for Biotechnology Information, National Library of Medicine, Bethesda, MD) ensuring products were free of primer dimers.

Total RNA was extracted from the dissected and isolated forebrain of each adult fish using homogenization with TriZol (Ambion) according to manufacturer protocol. Concentration of extracted RNA was obtained using NanoDrop 2000 (Thermo Scientific). Synthesis of cDNA was accomplished by reverse transcription of total RNA. From control and lesion specimens 500ng of total RNA were reverse transcribed using the Quantitect reverse transcription kit (Qiagen). Quality and purity of cDNAs was confirmed by Agarose gel Electrophoresis. In order to assay transcripts of genes of interests by qPCR, the following conditions were used: 95°C for 30s, followed by 40 cycles of 95°C for 5s and 59°C for 5s, then a melt curve progressing from 65°C to 95°C, at 5s per 0.5°C increase. Two reference genes were used for each qPCR either ef1a, ywhaz or rpl13a. Data were analyzed using CFXManager (Bio-Rad) and compiled using GraphPad PRISM.

### Statistical analyses

Statistical comparison of two groups (lesion and controls) for GFP cell counting values and qRT-PCR results was conducted using an unpaired t-test using GraphPad PRISM. An alpha-value of 0.05 was defined as statistically significant. For * p ≤ 0.05 and n.s. = not significant p > 0.05. Error bars represent standard error on the mean (SEM). Cell counts were performed on a minimum of two individuals in a blinded fashion to eliminate researcher bias.

## Results

### *dlx1a, dlx2a, dlx5a* and *dlx6a* are expressed in the adult zebrafish brain

Expression of *dlx* genes in the zebrafish brain has been reported during development (6,7,13,22), but information on the expression of such genes in adult zebrafish was still lacking. To determine if *dlx1a, dlx2a, dlx5a* and *dlx6a* are expressed in the adult zebrafish brain, we performed ISH assays in adults ranging from 3mpf to 18mpf.

Consistent with previous observations in embryos and larvae, the expression of all four *dlx* paralogs was abundant in ventral regions of the forebrain. Transcripts of *dlx1a, dlx2a, dlx5a* and *dlx6a* were present especially in the dorsal, ventral and postcommissural nucleus of the ventral telencephalic area (Vd, Vv and Vp) (Fig 1B - B’’’). For *dlx2a* and *dlx5a*, expression was also found within the central nucleus of the ventral telencephalic area (Vc). The anterior part of the parvocellular preoptic nucleus (PPa) was observed to be one of the regions with the most abundant expression of all four *dlx* genes in the adult zebrafish brain (Fig 1C-C’’’). In the midbrain, the caudal and dorsal zones of the periventricular hypothalamus (Hc and Hd) revealed abundant expression of all four *dlx* genes (Fig 1D-D’’’). The expression of *dlx* genes was consistent and similar among the four different paralogs as well as among different stages of the adult zebrafish, ranging from 3mpf to 18mpf (N=8 for each gene), therefore not presenting an age-dependent variation during adulthood.

**Fig 1.**
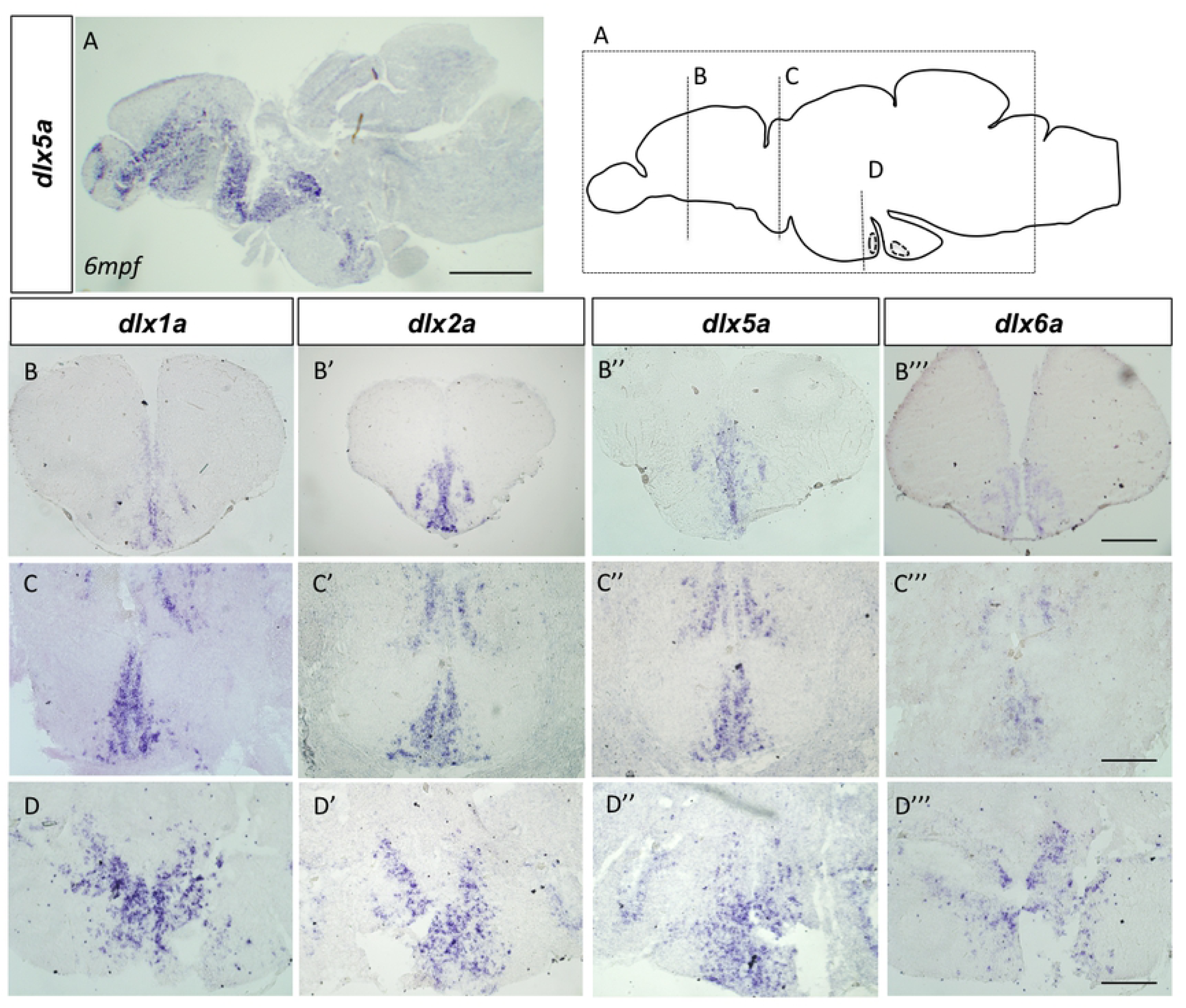
*dlx1a, dlx2a, dlx5a* and *dlx6a* are expressed in the adult zebrafish brain. Schematic representation of sections depicted in the top right panel. *In situ* hybridization shows *dlx* expression in cryosections of the adult zebrafish brain. Sagittal section showing *dlx5a* expression in a 1ypf fish (A). Transverse sections showing expression throughout areas of the forebrain, midbrain and hindbrain of *dlx1a* (B-D), *dlx2a* (B’-D’), *dlx5a* (B’’-D’’) and *dlx6a* (B’’’-D’’’) in 1 ypf zebrafish (N=6 for each *dlx* gene). Scale bar (A): 1mm; (B-D’’’) : 400µm

Our observations also reveal that all four *dlx* paralogs are expressed in almost all niches which present constitutive proliferation in the adult zebrafish brain, providing the first suggestions that these genes may participate in constitutive proliferation (Supp Fig1). Although the expression of *dlx1a, dlx2a, dlx5a* and *dlx6a* was observed to be highly overlapping in the adult zebrafish brain, we cannot conclude from our results if the different *dlx* paralogs are co-expressed within the same individual cells.

### GABAergic neurons identity for *dlx*-expressing cells in the adult zebrafish brain

To determine the identity of cells expressing *dlx1a, dlx2a, dlx5a* and *dlx6a* in the adult zebrafish brain, we performed co-localization analyses using double fluorescence *in situ* hybridization (ISH) and double immunohistochemistry assays. Based on the relationship between *dlx* and *gad1* expression as well as on the regulatory roles for *dlx* genes in GABAergic neuron development, we first wanted to analyze if *dlx*-expressing cells could have a GABAergic interneuron identity. Double fluorescence ISH assays were performed combining a mRNA probe recognizing one of the four *dlx* paralogs with *gad65*, a gene encoding an enzyme that catalyzes the decarboxylation of glutamate to GABA.

Widespread co-expression of *dlx* genes and *gad65* was observed throughout the adult zebrafish forebrain. The majority of *dlx2* and *dlx5a*-expressing cells co-expressed *gad65* in the medial zone of the dorsal telencephalic area (Dm), dorsal nucleus of ventral telencephalic area (Vd), postcommissural nucleus of ventral telencephalic area (Vp), parvocellular preoptic nucleus, anterior part (PPa) and posterior part of parvocellular preoptic nucleus (PPp) (Fig 2I-L) (N=4 for *dlx2a* and *dlx5a*). Similar observations were obtained for *dlx1a* and dlx6a (data not shown). In the midbrain, co-expression of *gad65* and *dlx1a, dlx2a, dlx5a* and *dlx6a* was less prevalent compared to areas of the forebrain. In the ventral zone of the periventricular hypothalamus (Hv) and dorsal zone of the periventricular hypothalamus (Hd), only a small portion of *dlx*-expressing cells presented a GABAergic interneuron identity (Supplementary Fig 2).

**Fig 2.**
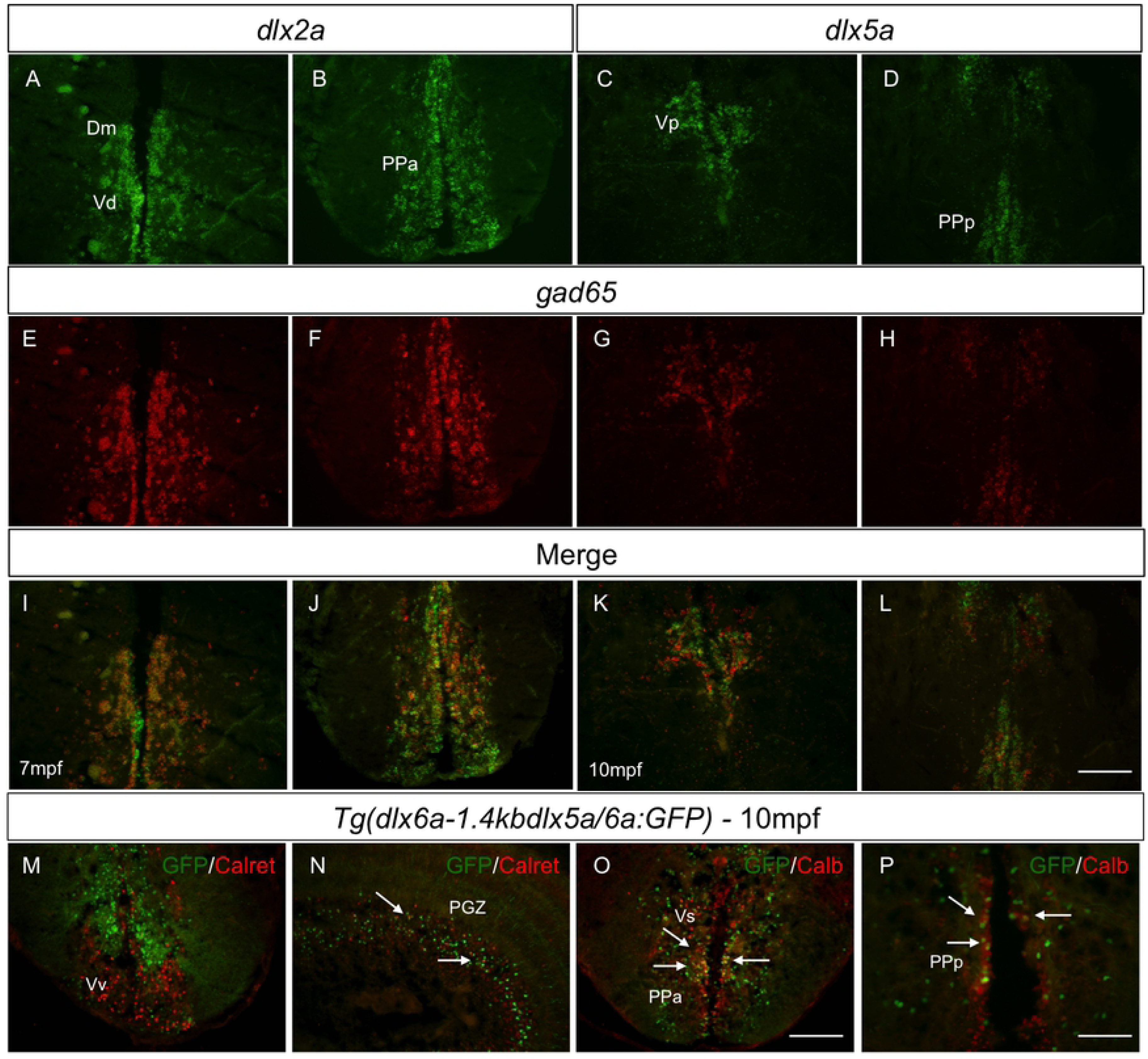
Co-expression of *dlx* paralogs with markers of GABAergic neurons in adult zebrafish. Double fluorescence ISH of transverse sections of the forebrain showing expression of *dlx2a* (A-B) and *dlx5a* (C-D) in green and expression of *gad65* in red (E-H). Anatomical parts indicated. Merged images showing co-localization of *dlx* and *gad65* in yellow [I-L] (N=4 for *dlx2a* and *dlx5a*). Double IHC with *Tg(dlx6a-1.4kbdlx5a/6a:GFP)* and Calretinin or Calbindin shows co-localization, indicated by white arrows [N-P] (N=6 for Calbindin and Calretinin). Merged images were created with ImageJ(32) software. Calret.: Calretinin and Calb.: Calbindin. Dm.: medial zone of dorsal telencephalic area; PGZ.: periventricular gray zone; PPa.: anterior part of parvocellular preoptic nucleus; PPp.: posterior part of parvocellular preoptic nucleus; V.: ventral telencephalic area; Vd.: dorsal nucleus of V.; Vp.: parvocellular nucleus of V.;Vs.: supracommissural nucleus of V.; Vv.: ventral nucleus of V. Scale bar: 400µm

As there is absence of good antibodies that recognize the *Dlx* proteins in zebrafish, the *Tg(dlx6a-1.4kbdlx5a/6a:GFP)* line allowed us to better investigate co-localization and quantifications of *dlx5a/6a*-expressing cells Even though all GABAergic neurons present inhibitory functions, these neurons can be morphologically, electrically and chemically heterogeneous, and there are several subtypes of GABAergic neurons in the CNS. Using *Tg(dlx6a-1.4kbdlx5a/6a:GFP)* adult fish, we investigated if *dlx5a/6a*-expressing cells could be specifically labeled to some of the subtypes of GABAergic interneurons, namely: calbindin and calretinin. Similar to observations at developmental stages, our analyses indicate that, with a few exceptions, the majority of *dlx5a/6a*-expressing cells do not co-localize with these specific subtype markers (Fig. 16). Co-labeling with calretinin has shown very little if any co-localization with GFP. Only a few *dlx5a/6a*-expressing cells appear to be calretinin neurons in the periventricular gray zone (PGZ) (Fig 2M-N). In fact, within the PGZ area, a few *dlx5a/6a*-expressing cells have also shown a calbindin identity.

Interestingly and in contrast to observations at developmental stages, in the adult zebrafish brain, we observed many *dlx5a/6a*-expressing cells co-localizing with calbindin within the supracommissural nucleus of the ventral telencephalic area (Vs) and within the anterior and posterior part of parvocellular preoptic nucleus (PPa and PPp) in the diencephalon (Fig 2O-P). A few GFP positive cells also co-express calbindin within the dorsal and ventral nucleus of the ventral telencephalic area (Vd and Vv). No co-localization was observed in areas of the hypothalamus.

### Neural stem cells, but not proliferating or glial cells, express *dlx5a/6a* in areas of the adult zebrafish brain

As the *dlx* genes might be implicated in promoting neuronal proliferation (25) (26), we sought to investigate if neural stem cells (NSCs) could also express *dlx* genes. Our results indicate that cells expressing *dlx5a/6a* co-localized with the sex-determining 2 (Sox2) marker in some areas of the adult zebrafish brain, while cells expressing GFP and Sox2 were adjacent in others. The following areas of the forebrain presented a small percentage of co-localization of the two markers: the medial dorsal telencephalic area (Dm) in the dorsal and ventral nucleus of the ventral telencephalic area (Vd and Vv). Within the supracommissural nucleus of the ventral telencephalic area (Vs), we observed a higher percentage of co-localization than in the domains of the telencephalon as mentioned before (Fig 3).

**Fig 3.**
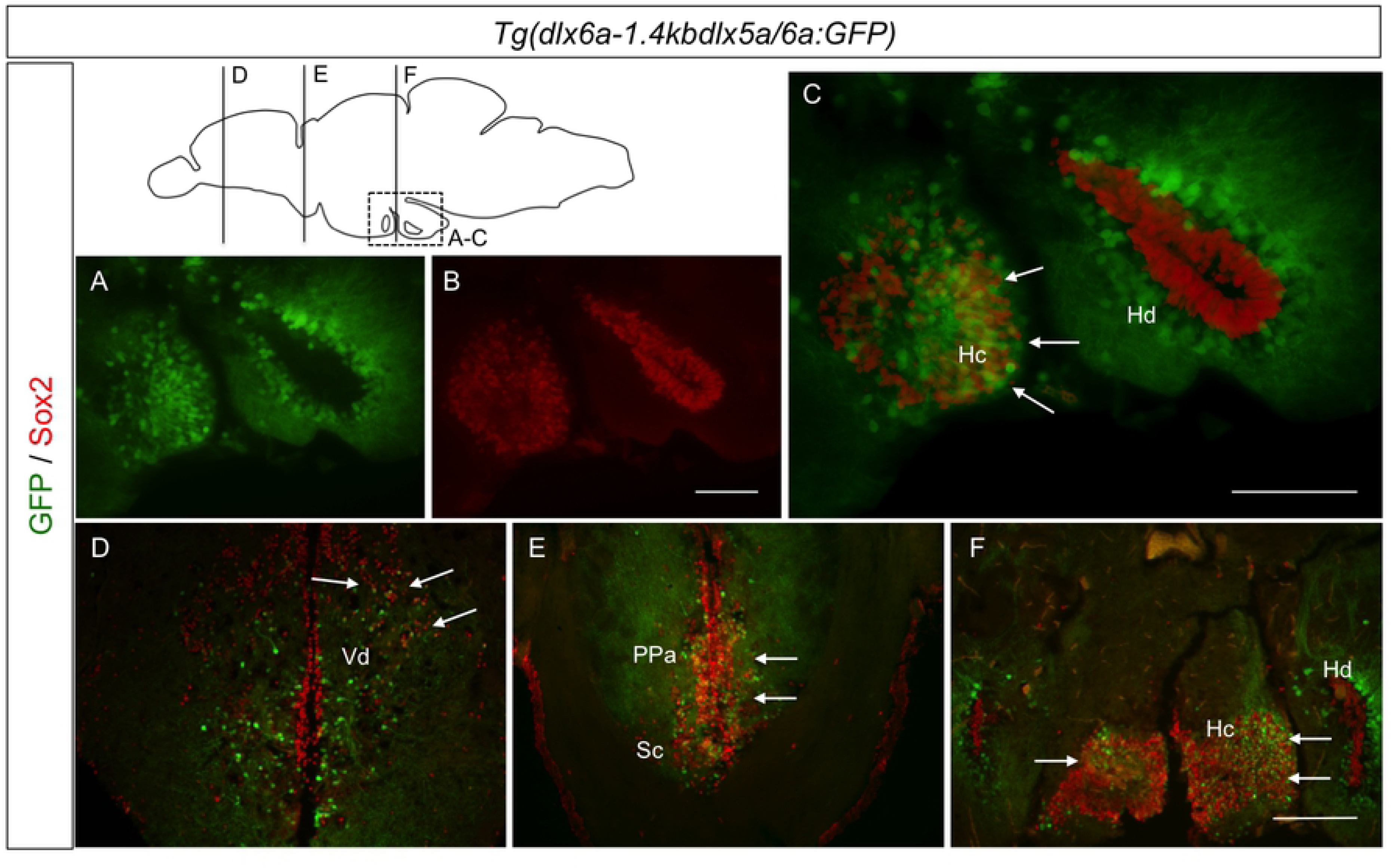
Co-localization of GFP and Sox2 in the *Tg(dlx6a-1.4kbdlx5a/6a:GFP)* adult zebrafish brain. Double IHC with *Tg(dlx6a-1.4kbdlx5a/6a:GFP)* and Sox2 shows co-localization, indicated by white arrows, in merged images of GFP and Sox2 (C-F) (N=8). Merged images were created with ImageJ(32) software. Hc.: caudal hypothalamus; Hd.: dorsal zone of periventricular hypothalamus; PPa.: anterior part of parvocellular preoptic nucleus; Sc.: suprachiasmatic nucleus; Vd.: dorsal nucleus of V. Scale bar: 400µm

The anterior part of the parvocellular preoptic nucleus (PPa), within the diencephalon, was one of the areas with high co-localization of *dlx5a/6a*-expressing cells and Sox2 expression. The more rostral portions of the dorsal and ventral hypothalamus presented some co-localization of GFP and Sox2, and the most caudal portions of the hypothalamus also seemed to reveal a high percentage of GFP and Sox2 co-localization (Fig 3C, E-F).

These data suggest that, in some areas of the adult zebrafish brain, a number of *dlx5a/6a*-expressing cells present a neural stem cell identity, especially in the PPa and the hypothalamus, two areas where expression of all four *dlx* paralogs is very abundant. Giving the overlapping expression within these areas, these *dlx5a/6a*-expressing cells might have a role in promoting neural proliferation during adulthood in the zebrafish brain. In all other areas of the adult zebrafish not mentioned before, the great majority of *dlx5a/6a*-expressing cells did not co-localize with Sox2 (data not shown). The adjacent expression of *dlx5a/6a* to Sox2, clearly observed in Hd for example (Fig 3C), also suggest that many *dlx*-expressing cells have already reached a more differentiated state.

We also examined if *Tg(dlx6a-1.4kbdlx5a/6a:GFP)*-expressing cells could represent either glia populations or proliferating cells. We observed rare, if any, co-localization of GFP with glial fibrillary acidic protein (GFAP) or with the proliferating cell nuclear antigen (PCNA) (Fig 4A-H), giving indications that in the adult zebrafish brain, *dlx5a/6a*-expressing cells might not have a proliferating or glial cell identity.

**Fig 4.**
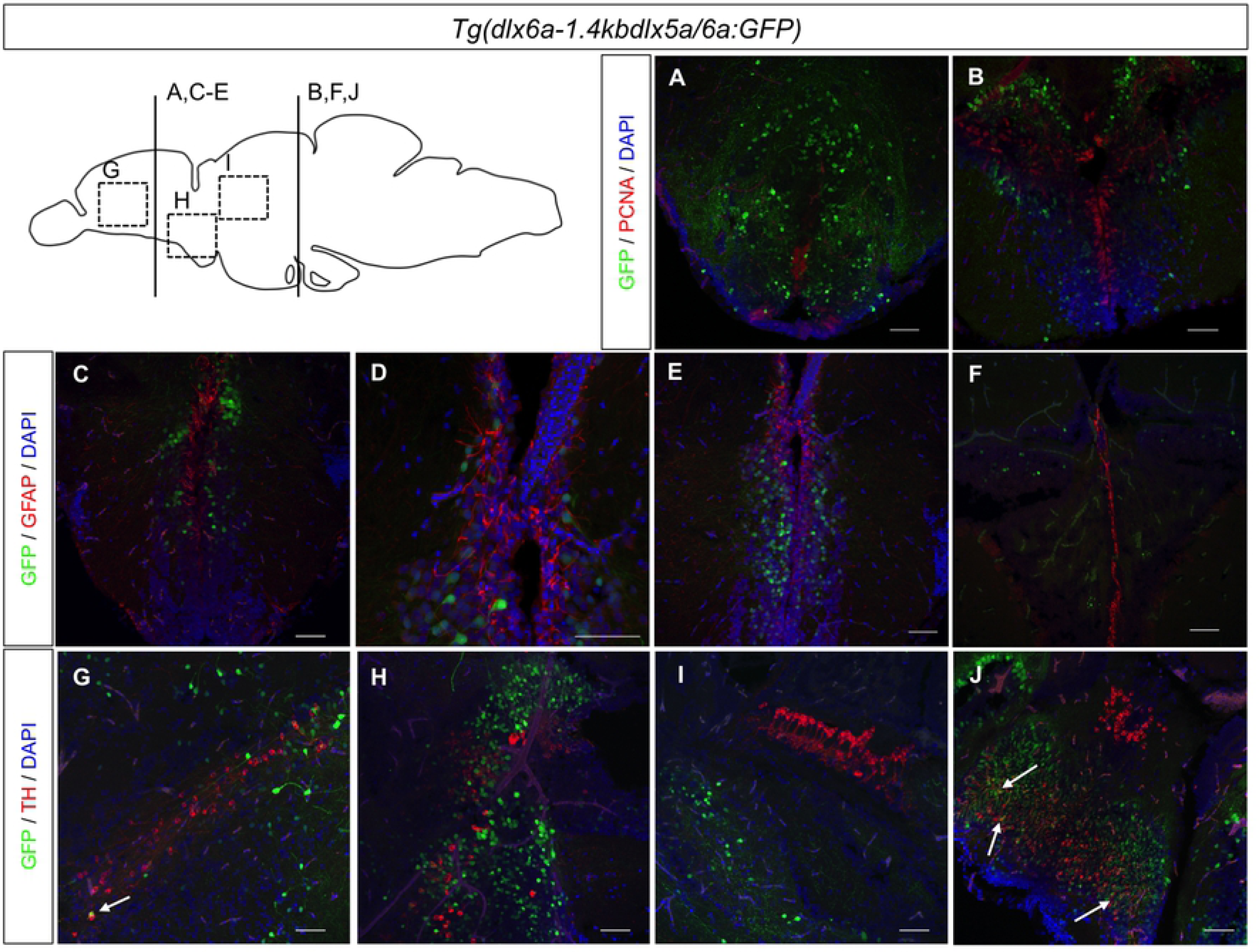
Immunohistochemical labeling of PCNA, GFAP and TH in *Tg(dlx6a-1.4kbdlx5a/6a:GFP)* adult zebrafish. Double IHC with *Tg(dlx6a-1.4kbdlx5a/6a:GFP)* in combination with either PCNA, GFAP or TH. Labeling of GFP with PCNA (A-D) and GFAP (E-H), shows no co-localization of the two markers with GFP (N=6). Labeling of GFP and TH (I-L) shows a few co-localizations of the two markers, indicated by white arrows. Merged images created with NIS-Elements Advanced Research Software. Scale bar: 200µm

A few studies have suggested a possible role for *dlx* genes in dopaminergic subtype specification and regulation (27) (28). Additionally, some evidence indicates co-expression of markers for GABAergic neurons and tyrosine hydroxylase (TH), an enzyme that catalyzes the first reaction in dopamine biosynthesis(29). We sought to investigate if, in the adult *Tg(dlx6a-1.4kbdlx5a/6a:GFP)* zebrafish brain, there was co-localization of GFP and TH. Once again, results show rare instances of a few single cells co-expressing GFP and TH in the ventral telencephalic area and in the caudal zone of the hypothalamus (Hc) (Fig 4I-L).

Taken together, these co-localization observations reveal that in the adult zebrafish brain, the majority of *dlx*-expressing cells seem to have a GABAergic neuronal identity. The results are summarized on Table 1.

### *dlx5a* and *dlx5a/6a* are up-regulated during brain regeneration following stab wound lesion

The *dlx* genes are required during development for proper establishment of neuronal populations in the central nervous system. The zebrafish brain presents high levels of adult neurogenesis and regeneration, as previously mentioned. Therefore, we explored a possible participation of *dlx* paralogs in adult brain regeneration. In order to address this, we have used mechanical stab lesions in the telencephalon, an area with both high rates of constitutive proliferation and high expression of all four *dlx* genes.

We analyzed the expression of *dlx1a, dlx2a, dlx5a* and *dlx6a* by *in situ* hybridization at 7 days-post-lesion(dpl) (3mpf to 1ypf). This time point presents a very important stage of the regeneration response and the peak of constitutive proliferation after a lesion (17). The spatial distribution of *dlx1a, dlx2a, dlx5a* and *dlx6a* transcripts remained similar during the regeneration response, with expression concentrated in the dorsal (Vd) and ventral (Vv) nucleus of the ventral telencephalon in the sections analysed (Fig 5F-I).

**Fig 5.**
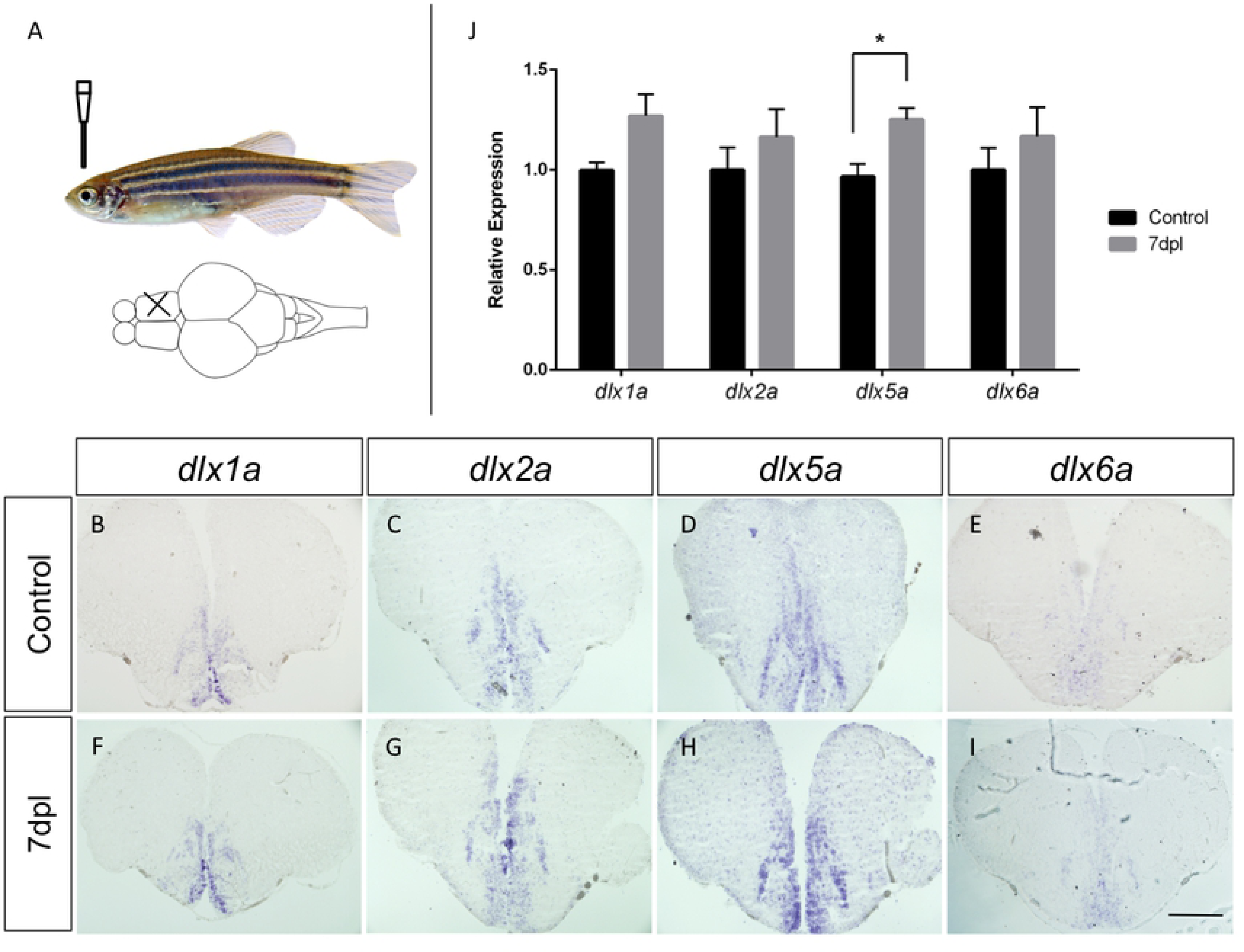
Expression of *dlx1a, dlx2a, dlx5a* and *dlx6a* post-lesion in adult zebrafish. Top left panel (A) shows the location of the mechanical lesion. Expression of the four *dlx* paralogs in controls (B-E) and lesioned brains (F-I). A slight up-regulation of *dlx2a* and *dlx5a* was apparent compared to controls (G and H). N=6 for each gene (2 biological replicates in 3 different experiments). RT-qPCR analyses with RNA extracted from the telencephalon of regenerating brains at 7dpl and control specimens (J). No significant changes in expression levels of *dlx1a, dlx2a* and *dlx6a* were observed (N=7 for each gene each). A significant increase in *dlx5a* expression was observed at 7dpl (*Student’s t-test*, n=7, p=0.008). Scale bar: 400 μM

A slight up-regulation of *dlx5a* at 7dpl was suggested based on the intensity of the ISH signal (Fig 5H). At this time point, expression of *dlx5a* consistently presented slight increases and a more widespread expression pattern within the dorsal telencephalic area. Overall, expression of all four paralogs is very weak in the dorsal telencephalon and ventricular zone of the telencephalon. This increase was verified by experimental repetition and biologic replicates (n= 6 for each *dlx* gene).

We did not observe apparent changes in the expression of *dlx1a, dlx2a* or *dlx6a* at 7dpl (Fig 5J), nor in the expression patterns of *dlx1a, dlx2a, dlx5a* or *dlx6a* at the site of injury where the needle was inserted in the ventricule of telencephalon at 7dpl.

Possible changes in expression levels of *dlx1a, dlx2a, dlx5a* and *dlx6a* were further quantified by qRT-PCR at 7dpl. Slight increases in *dlx1a, dlx2a* and *dlx6a* transcripts were seen but did not reach statistical significance at 7dpl. However, there was a significant increase in *dlx5a* expression in the telencephalon of lesioned adult zebrafish at 7dpl (Fig.5 J).

Changes in *dlx* expression during brain regeneration was further examined using the *Tg(dlx6a-1.4kbdlx5a/6a:GFP)* reporter line and direct counting of GFP positive cells. The dorsal and ventral areas of the ventral telencephalon is where constitutive proliferation takes place and are also the regions were an increase seemed more visible, therefore this area was selected for quantification (Fig.6). Cell counting revealed an increase in the number of GFP-expressing cells in the ventral portions of the telencephalon of regenerating brains of adults (9mpf) at 3 dpl (average 266 [180-384] vs. 233 [99-396] in controls, N=6) and 7pl (average 344 [244-408] vs. 247 [198-302] in controls, N=6). At 3 dpl, this increase was not significant (*Student’s t-test*, p=0.613 Fig. 25.F), while at 7dpl this number reached statistical significance (*Student’s t-test*, p=0.008 Fig. 6.F).

**Fig 6.**
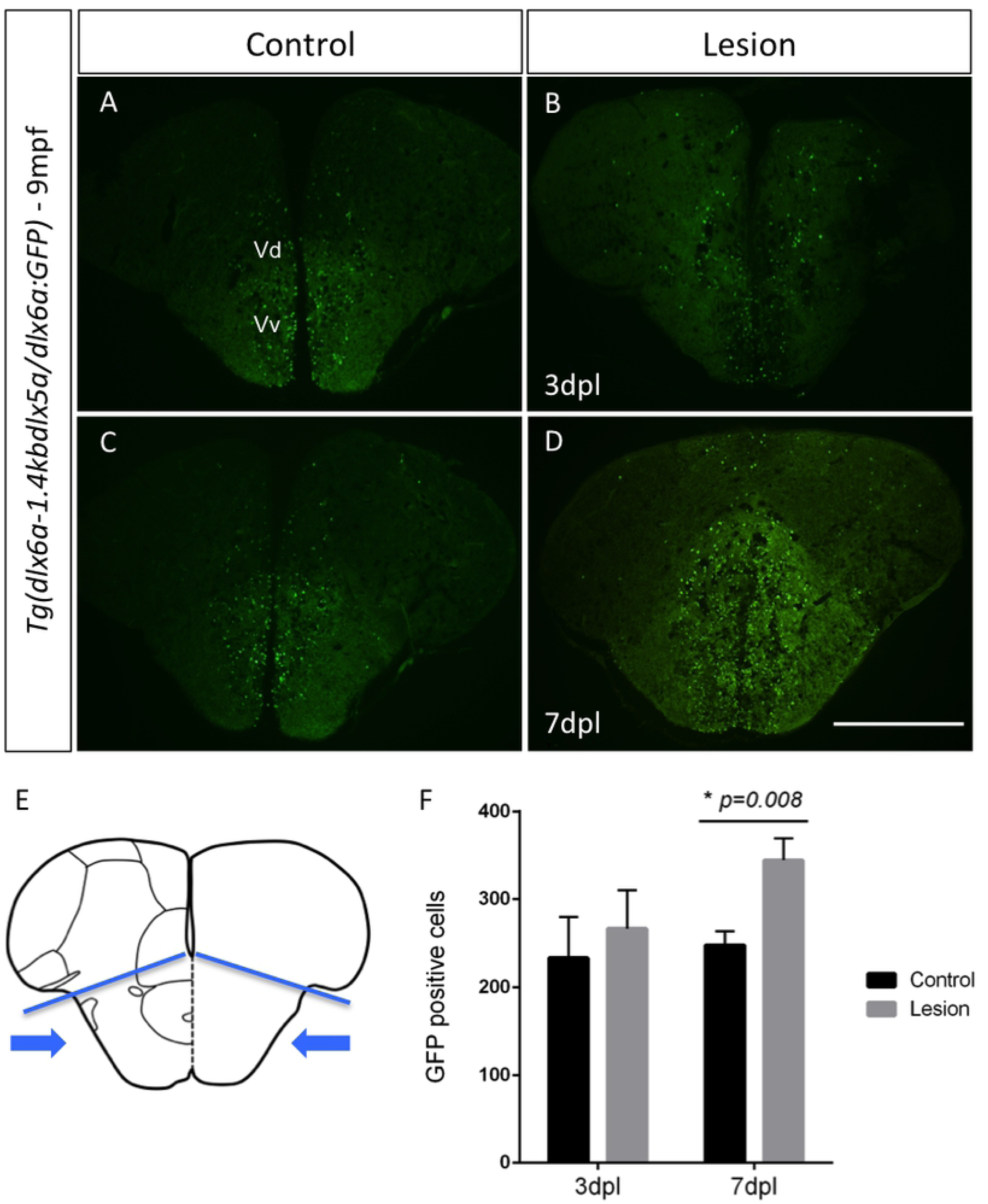
GFP labeling in *Tg(dlx6a-1.4kbdlx5a/6a:GFP)* adult zebrafish at 7 days post-lesion and cell quantification. Expression of GFP in *Tg(dlx6a-1.4kbdlx5a/6a:GFP)* determined with a GFP antibody in controls (A and C) and regenerating brains at 3 dpl (B) and 7dpl (D). Schematic representation of the telencephalon where lesion is inflicted and areas used for cell counting of all GFP-positive cells (E) (area indicated by blue arrows bellow blue lines). Quantification of GFP+ cells in the regenerating brain at 3dpl and 7dpl (P=0.008, N=6) (F). Scale bar: 400 μm

These data suggest that, at 7 dpl, a time when the regeneration response is pronounced in the adult zebrafish brain, the *dlx5a/6a* bigene, possibly the *dlx5a* gene, may participate in and reflect an increased proliferation within the ventral area of the ventricule of the telencephalon.

## Discussion

The observations presented here expand our current understanding of *dlx* function from the context of development to adulthood in zebrafish. Here, we report the expression of *dlx* genes in the adult zebrafish brain, characterize the GABAergic and NSCs identity of cells expressing *dlx* in adults, and verify the expression of these genes, particularly of *dlx5a* and *dlx5a/6a*, during regeneration in a post-injury response.

The first observations of *dlx1a, dlx2a* and *dlx5a* expression in the zebrafish developing brain indicated the onset of expression at around 13h hours-post-fertilization (6) (7). The expression of the four paralogs *dlx1a, dlx2a, dlx5a* and *dlx6a*, is present throughout embryonic and larval stages as demonstrated by others (11) (22) (30). In adults, we observed that *dlx1a, dlx2a, dlx5a* and *dlx6a* are expressed similarly among the four different paralogs and independently of adult stages ranging from 3mpf to 18mpf.

In the forebrain, the domains of *dlx1a, dlx2a, dlx5a* and *dlx6a* expression, namely the dorsal, ventral and parvocellular nucleus of ventral telencephalic area and diencephalon, remained consistent with observations made during embryonic development. Transcripts of *dlx2a* and *dlx5a* were also observed within the central nucleus of the ventral telencephalic area (Vc). In fact, in many areas, *dlx2a* and *dlx5a* expression was seemingly more abundant than *dlx1a* and *dlx6a*. This was not unexpected as, in embryos, the intensity of the *dlx5a* ISH signal was comparatively more uniform and stronger than that of *dlx6a* (7). However, we do not rule out the possibility of the results obtained here being due to less effective hybridization of mRNA probes utilized for *dlx1a* and *dlx6a* in ISH assays. In contrast to what was observed in the diencephalon of embryos, the anterior and posterior part of parvocellular preoptic nucleus were regions with abundant expression of all four *dlx* genes in the adult zebrafish brain. Yet, other domains of the diencephalon such as the hypothalamus, caudal and dorsal hypothalamus, presented abundant expression of *dlx1a, dlx2a, dlx5a* and *dlx6a* genes. The spatiotemporal expression of *dlx* genes in the adult brain could be indicative of multiple roles, ranging from cell fate determination to neurogenesis. In fact, many of the adult zebrafish brain regions where *dlx1a, dlx2a, dlx5a* and *dlx6a* are expressed consist of neurogenic zones (sup fig) (21).

Given previous observations of the participation of *dlx* genes in GABAergic interneuron specification (2) (8), we expected that in the adult zebrafish brain, many *dlx*-expressing cells would have a GABAergic neuronal identity. Indeed, our observations revealed that, in telencephalic regions, there is a high overlapping expression of *dlx* and *gad65* transcripts, indicating that the great majority of *dlx*-expressing cells consist of GABAergic neurons in these areas. In the diencephalon, however, at the ventral and dorsal zone of the periventricular hypothalamus (Hd and Hv), only a small portion of *dlx*-expressing cells presented a GABAergic interneuron identity (sup figure). This might indicate that, in these areas, those cells are in the cell cycle and have not yet acquired the GABAergic phenotype, or that *dlx*-expressing cells will give rise to different identities.

The calcium binding proteins calretinin and calbindin are expressed in GABAergic and glutamatergic cortical neurons (31). *Dlx* enhancers in mice have been shown to be highly active in some of the major subtypes of GABAergic interneurons (32). Interestingly, during the early development of zebrafish, the comparison of GFP expression in *Tg(dlx6a-1.4kbdlx5a/6a:GFP)* embryos with markers for GABAergic subtypes, revealed that a vast majority of GFP-positive cells within the telencephalon and diencephalon of 3 dpf embryos do not co-localize with any of these markers (22). Similar to what was observed during development, our results indicate that the majority of *dlx5a/6a*-expressing cells do not co-localize with these specific subtypes in the adult zebrafish brain, with some exceptions. However, the anterior and posterior part of parvocellular preoptic nucleus as well as the supracommissural nucleus of ventral telencephalic area presented cluster of cells with high co-localization of GFP and Calbindin, suggesting that *dlx5a/6a* is highly active in Calbindin interneurons in these regions.

Apart from their known role in GABAergic neurons specification, the *dlx* genes can be considered pro-neural transcription factors known to promote neural proliferation (26). Our results revealed that cells expressing *dlx5a/6a* genes do not seem to have a glial or proliferating cell identity in the adult brain as no co-localization was observed with GFAP or PCNA markers. Interestingly, in some areas of the forebrain and midbrain, particularly in the anterior part of parvocellular preoptic nucleus, the supracommissural nucleus of ventral telencephalic area and caudal hypothalamus we observed high overlapping co-localization of the neural stem cell marker Sox2 and GFP in the brain of *Tg(dlx6a-1.4kbdlx5a/6a:GFP)* adults. These observations suggest a role for *dlx* genes in the maintenance of neural pluripotency or in promoting neural proliferation in the adult brain.

We frequently observed marker co-localization in the telencephalon and diencephalon that differed from that observed within the hypothalamus domains of the adult zebrafish brain. The hypothalamus is involved in the regulation of body temperature and reproduction and can be considered a central interface in which neuronal, hormonal and vascular systems are connected (33). Other transcription factors have been implicated in neuronal specification within the hypothalamus of the zebrafish (34). Certain areas of the hypothalamus, especially in the caudal and dorsal hypothalamus (Hc and Hd, respectively), presented very little co-localization of *dlx* transcripts and *gad65* (Suppl. Fig.2). It was also within the Hc that the highest overlapping expression levels of *dlx* genes and Sox2 were observed, as well as a few occasions of co-localization of TH and GFP with *Tg(dlx6a-1.4kbdlx5a/6a:GFP)* adult zebrafish. While co-localization of *Sox2* and *dlx5a/6a* was abundant in the Hc, the expression of these genes was observed in adjacent patterns in the Hd. This suggests that while in some areas of the hypothalamus these genes may be involved with the reprograming of cells to become mature neurons, in other areas of the hypothalamus, *dlx* transcripts may be present in recently formed mature neurons that do not have a GABAergic identity.

Due to the intense reactive proliferation in the brain during regeneration (16,17), we expected to see changes in the patterns of expression of *dlx1a, dlx2a, dlx5a* and *dlx6a*, as these genes take part in important developmental events and neuronal specification. We observed a slightly stronger ISH staining expression of *dlx2a* and *dlx5a* in the telencephalon of regenerating brain at 7dpl. This time point is thought to represent a very important stage of the regeneration response and the peak of constitutive proliferation after mechanical lesion (24) (35). The expression of *dlx5a* appeared to be particularly stronger within the the dorsal and parvocellular nucleus of ventral telencephalic area, and this gene showed more widespread expression patterns within the dorsal telencephalic area. Although these increases were observed for *dlx2a* and *dlx5a*, in both cases, the presence of transcripts was not found adjacent or exactly at the location of injury at the ventricular zone.

Counting of GFP positive cells with the *Tg(dlx6a-1.4kbdlx5a/6a:GFP)* reporter line revealed a significant increase in *dlx5a/6a*-expressing cells in specific areas of the telencephalon at 7dpl. In this reporter line it is not fully known if the GFP reporter recapitulates the expression of *dlx5a* and *dlx6a* equally, additionally, there is an increased sensitivity of the GFP reporter and easier detection than mRNA transcripts with ISH (36). RNA quantification analyses of the telencephalon of lesioned adult zebrafish corroborate the results observed by ISH. Slight increases in *dlx1a, dlx2a* and *dlx6a* transcript levels were observed at 7dpl and statistically significant increases were obtained for *dlx5a*. These results suggest that *dlx* genes may participate in post-injury response. Thus, increased expression of these genes may participate in compensating for neuronal loss, specifically the loss of GABAergic neurons.

The events subsequent to a traumatic lesion in the CNS can lead to an increase in neurogenesis depending mainly on three aspects: the severity of the lesion, the site of the trauma, and the competency of the progenitor cells (21,37,38). The adult mammalian brain harbours neural precursor cells (NSCs), which are a potential source of neurons for repairing injured brain tissue. Recent studies show that the telencephalic ventricular zone in the adult zebrafish brain, where the dorsal and ventral nucleus of ventral telencephalic area are located, contain NPCs that share characteristics with the NPCs in the mammalian SVZ (20) (39) (40) (41). The results obtained here reveal potential roles for the *dlx* genes in a regeneration response towards reappearing injured brain tissue.

Given the important roles already described for *dlx* genes in the CNS during development, the work presented here expands our knowledge of *dlx* genes function to the context of adulthood. Understanding the role of transcription factors in the adult CNS as well as the mechanisms involved in regeneration biology of the vertebrate CNS present great potentials for therapies, especially regarding human neurodegenerative disorders or acute neural injuries.

## Supplementary Information

**Supplementary Table 1.**
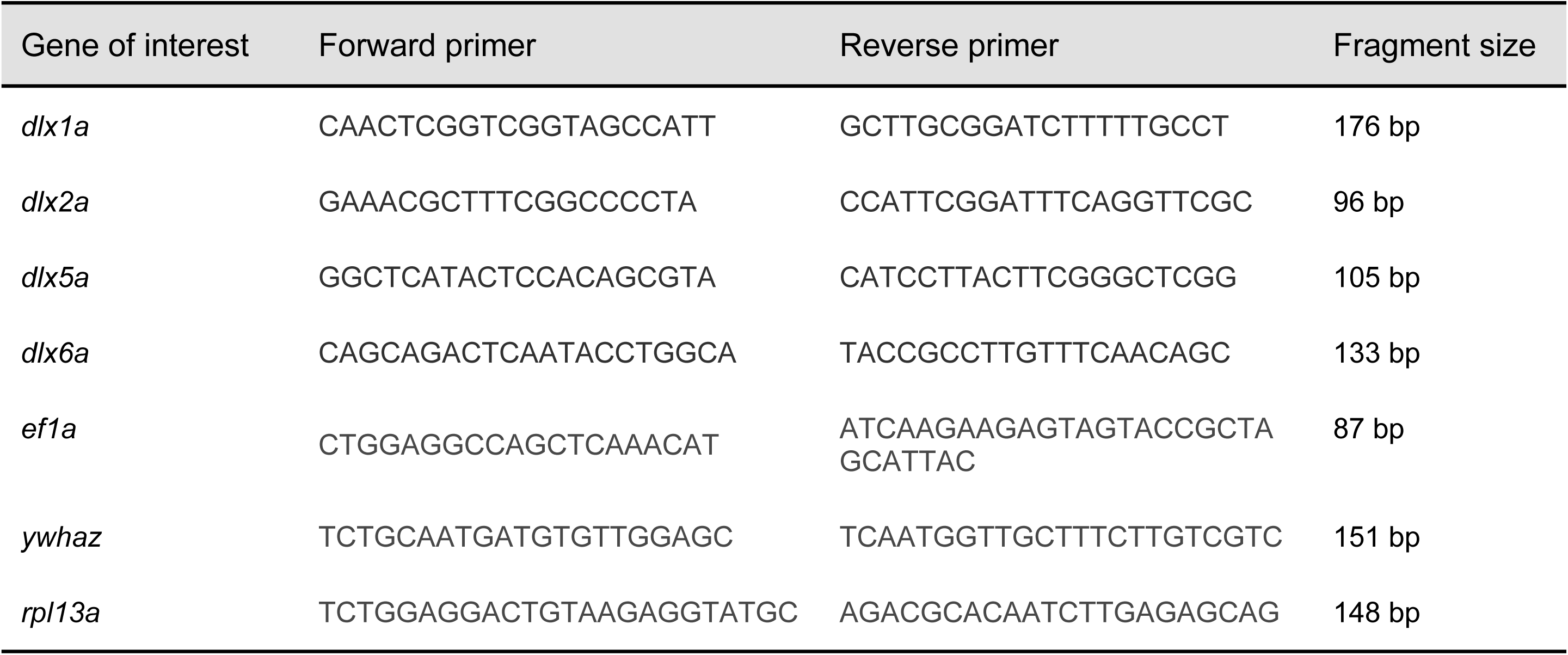
Primers used for qRT-PCR.

**Supplementary Figure 1. *dlx1a* and *dlx5a* expression and comparison with neurogenic areas in the adult zebrafish brain.**

Upper panel shows drawing of an adult brain sagittal section depicting neurogenic areas and zones with constitutive proliferation (adapted from Kizil C. *et al.*, 2011). [A-B] shows expression of *dlx1a* and *dlx5a* verified with ISH. Arrows indicate the main areas where expression of dlx genes matches areas with constitutive proliferation. These areas are: olfactory bulb (OB), ventral nucleus of ventral telencephalic area (Vv), parvocellular preoptic nucleus, anterior part (PPa), posterial zone of dorsal telencephalic area (Dp), periventricular nucleus of posterior tuberculum (TPp) and caudal zone of periventricular hypothalamus (Hc) and dorsal zone of periventricular hypothalamus (Hd).

Scale bar: 1mm

**Supplementary Figure 2. Co-expression of *dlx2a* and *dlx5a* with *gad65* in the adult zebrafish forebrain.**

Double fluorescence *in situ* hybridization with transverse sections of the forebrain with [A-D] expression of *dlx2a* and *dlx5a* in green along with anatomical parts indicated and [E-H] expression of *gad65* in red. [I-L] Co-localization of *dlx* and gad65 in yellow. Merged images were created with ImageJ(32) software. (N=4 for *dlx2a* and *dlx5a;* N=3 for *dlx1a* and *dlx6a*).

Scale bar: 50µm

## Acknowledgements

We would like to thank Yuchen Luo for contributions to some of the experiments, Vishal Saxena and Gary Hatch for technical support and also Dr. Marie-Andree Akimenko for discussions and suggestions.

## Author Contributions

Conceptualization of the study and design of experiments: HWM and ME. Experiments and data analysis: HWM, MT and TD. Writing of original draft: HWM. Reviewing and editing: MT, TD and ME. Funding acquisition and resources: ME.

